# Tau accumulation activates STAT1 triggering memory deficits via suppressing NMDA receptor expression

**DOI:** 10.1101/437814

**Authors:** Xiao-Guang Li, Xiao-Yue Hong, Ya-li Wang, Shu-Juan Zhang, Jun-Fei Zhang, Xia-Chun Li, Yan-Chao Liu, Dong-Shen Sun, Qiong Feng, Jin-Wang Ye, Yuan Gao, Dan Ke, Qun Wang, Hong-lian Li, Keqiang Ye, Gong-Ping Liu, Jian-Zhi Wang

**Affiliations:** Department of Pathophysiology, School of Basic Medicine and the Collaborative Innovation Center for Brain Science, Key Laboratory of Ministry of Education of China and Hubei Province for Neurological Disorders, Tongji Medical College, Huazhong University of Science and Technology, Wuhan 430030, China; Department of Pathology and Laboratory Medicine, Emory University School of Medicine, Atlanta, GA 30322, USA; Co-innovation Center of Neuroregeneration, Nantong University, Nantong, JS 226001, China; Clinic Center of Human Gene Research, Union Hospital, Tongji Medical College, Huazhong University of Science and Technology, Wuhan 430030, China; Department of Physiology and Neurobiology, Key Laboratory for the Brain Research of Henan Province, Xinxiang Medical University, Xinxiang, PR 453000, China

**Keywords:** Tau, STAT1, synapse, N-methyl-D-aspartate receptors, memory

## Abstract

Intracellular tau accumulation forming neurofibrillary tangles is hallmark pathology of Alzheimer's disease (AD), but how tau accumulation induces synapse impairment is elusive. By overexpressing human full-length wildtype tau (termed hTau) to mimic tau abnormality as seen in the brain of sporadic AD patients, we found that hTau accumulation activated JAK2 to phosphorylate STAT1 (Signal Transducer and Activator of Transcription 1) at Tyr701 leading to STAT1 dimerization, nuclear translocation and its activation. STAT1 activation suppressed expression of N-methyl-D-aspartate receptors (NMDARs) through direct binding to the specific GAS element of GluN1, GluN2A and GluN2B promoters, while knockdown STAT1 by AAV-Cre in STAT1^flox/flox^ mice or expressing dominant negative Y701F-STAT1 efficiently rescued hTau-induced suppression of NMDARs expression with amelioration of synaptic functions and memory performance. These findings indicate that hTau accumulation impairs synaptic plasticity through JAK2/STAT1-induced suppression of NMDARs expression, revealing a novel mechanism for hTau-associated synapse and memory deficits.

## INTRODUCTION

Intracellular accumulation of tau forming neurofibrillary tangles is one of the two hallmarks in Alzheimer’s disease (AD), the most common neurodegenerative disorder in the elderly [1, 2]. Abnormal tau accumulation is positively correlated with neurodegeneration and memory deterioration [3, 4], and the total tau level in cerebrospinal fluids has an inverse correlation with memory score in AD patients [5, 6]. The axonal tau pathology in hippocampus is critical for the clinical presentation of dementia and may constitute an anatomical substrate of clinically verifiable memory dysfunctions [3]. The human tau transgenic mice recapitulate features of human tauopathies and cognitive deficits [7, 8]. Tau is essential for β-amyloid-induced synaptic toxicity [9], while tau knockout attenuates neuronal dysfunction and prevents behavioral deficits in transgenic mice expressing human amyloid precursor protein (APP) without altering high Aβ level in the brain [10, 11]. These clinical and laboratory evidence strongly suggest that tau abnormality plays a pivotal role in AD-like synapse and memory impairments.

As a cytoskeleton protein, the originally characterized function of tau is to promote microtubule assembly and maintain the stability of microtubules, which is essential for axonal transport [12, 13]. Tau hyperphosphorylation dissociates microtubules and thus disrupts axonal transport [14–18]. Recent studies suggest that tau phosphorylation is actively involved in regulating cell viability [19–21]. Normally, tau is largely located in the neuronal axons [22]. Upon hyperphosphorylation [23], tau is located into the dendritic spines where it interacts with the postsynaptic proteins and thus induces synaptic dysfunction [24, 25]. Intracellular accumulation of tau causes mitochondrial dysfunction and mitophagy deficits by increasing mitochondrial membrane potential [26, 27]. Tau accumulation also disrupts intracellular calcium signaling leading to activation of calcineurin and CREB dephosphorylation in primary neuron cultures [4]. These hypothesis-driven studies partially disclose the mechanisms underlying the toxic effects of tau. However, the molecular mechanism underlying hTau-induced synapse impairment is not fully understood.

In the present study, we employed a large scale screening approach to explore novel molecular mechanisms underlying tau toxicities. By using whole-genome mRNA chip and the transcription factor activation profiling array, we found that overexpressing hTau upregulated JAK2/STAT1 signaling, and simultaneous downregulating STAT1 by hippocampal infusion of AAV-Cre in STAT1^flox/flox^ mice or by overexpressing dominant negative STAT1 mutant mitigates the hTau-induced synaptic and memory deficits. We also found that STAT1 can directly bind to the specific GAS elements GluN1, GluN2A and GluN2B and thus suppress expression of the NMDARs, which reveals a novel mechanism underlying hTau-induced synapse impairment and memory deficit.

## METHODS AND MATERIALS

### Antibodies and reagents

The antibodies used in the present study were listed in the Supplementary Table 3. Itacitinib (JAK1 inhibitor, from MCE), TG-101348 (special JAK2 inhibitor, from MCE), JAK2 siRNA (sc-39099, from Santa Cruz), AG490 (JAK2 inhibitor, from Santa Cruz), SP600125 (the inhibitor of JNK1, from Santa Cruz) and FR180204 (the inhibitor of ERK1, from Santa Cruz) were purchased. Human pIRES-eGFP-hTau plasmid was a gift of Dr. Khalid Iqbal (New York State Institute for Basic Research in Developmental Disabilities, Staten Island, NY). WT-STAT1 and Y701F-STAT1 plasmids were gift of Dr. Xiao-Yuan Li (Institute of Biomedical Sciences, Academia Sinica, Taiwan).

### Animals

Male C57 mice were purchased from the animal center of Tongji Medical College, Huazhong University of Science and Technology. STAT1flox/flox (signal transducer and activator of transcription 1) mutant mice (B6; 129S-STAT1tm1Mam/Mmjax) and hTau transgenic mice (STOCK Mapttm1(EGFP) Klt Tg(MAPT)8cPdav/J) were purchased from Jackson lab. All mice were kept at 24 ± 2 °C on daily 12 h light-dark cycles with ad libitum access to food and water. All animal experiments were performed according to the ‘Policies on the Use of Animals and Humans in Neuroscience Research’ revised and approved by the Society for Neuroscience in 1995, and the Guidelines for the Care and Use of Laboratory Animals of the Ministry of Science and Technology of the People’s Republic of China, and the Institutional Animal Care and Use Committee at Tongji Medical College, Huazhong University of Science and Technology approved the study protocol.

### Stereotaxic brain injection

Adeno-associated virus-eGFP expressing human full-length tau or the control eGFP AAV (eGFP), AAV-Cre (Cre) and AAV-Y701F-STAT1 virus were purchased from OBio Biologic Technology Co., Ltd. The titer of the AAV-hTau or its control virus was 1.13×10^13^ v.g./ml, AAV Cre or AAV-Y701F-STAT1 was 5×10^12^ v.g./ml, and Syn-hTau-AAV was 1.99×10^13^ v.g./ml. The *in vivo* overexpression efficiency was measured by immunohistochemical staining and Western blotting after injection of the virus into the hippocampal CA3 of mice brains for 1 month. For brain injections, ~3 m-old C57 or STAT1flox/flox mice were positioned respectively in a stereotaxic instrument, then the virus was bilaterally injected into the hippocampal CA3 region (AP ±2.0, ML −1.5, DV −2.0) at a rate of 0.10 μl/min. The needle syringe was left in place for ~3 min before being withdrawn. The injection did not significantly increase the death rate or change the normal activity of the mice compared with the non-injected controls. The hippocampal CA3 region which infected with the virus was used for the biochemical measurements.

### Behavioral tests

Four weeks after brain infusion of the viral vectors, the spatial learning and memory were assessed by Morris water maze (MWM) test [28]. For spatial learning, mice were trained in water maze to find a hidden platform for 5 consecutive days, 4 trials per day with a 30 s interval from 14:00 to 20:00 pm. On each trial, the mice started from one of the four quadrants facing the wall of the pool and ended when the animal climbed on the platform. If the mice did not locate the platform within 60 s, they were guided to the platform. The swimming path and the time used to find the platform (latency) or pass through the previous platform quadrant were recorded each day by a video camera fixed to the ceiling, 1.5 m from the water surface. The camera was connected to a digital-tracking device attached to an IBM computer. The spatial memory was tested 1 day after the last training. The longer a mouse stayed in the previous platform-located quadrant, the better it scored the spatial memory.

The fear conditioning test was performed as the procedures established in our lab [29]. Briefly, the mouse was kept in the cage for 3 min to adapt to the environment before experiments, and then the mice received training by subjecting to 3 min unsignaled foot-shocks (one shock at the first min, three shocks at the second min and 8 shocks at the third min; 0.5 mA, 2-sec duration, and 1 min apart). The short-term memory (STM) and long-term memory (LTM) were tested respectively in 2 h and 24 h after the training by subjecting back into the conditioning chamber for 3 min and measuring the freezing time.

### Electrophysiological analysis

Horizontal brain slices (400 μm) containing the dorsal hippocampus were cut at 4-5 °C in artificial cerebrospinal fluid (aCSF) consisting of (in mM): 126 NaCl, 3 KCl, 1.25 NaH_2_PO_4_, 24 NaHCO_3_, 2 MgSO_4_, 2 CaCl_2_ and 10 glucose (pH 7.4; 305 mOsm), and saturated with carbogen (95% O_2_ and 5% CO_2_), using a Leica VT1000S vibratome (Milton Keynes, UK). Immediately after slicing sections were transferred and maintained in an interface chamber continuously perfused with aCSF. The slices were allowed to equilibrate at least for 30 min prior to recording at room temperature.

For extracellular recordings, slices were placed in the interface recording chamber at 32 °C and the perfusion rate was normally 3 ml/min, while maintaining a thin film of aCSF covering the slice to make sure applied substances could diffuse into the area recorded. Field potentials were amplified with Neurolog AC-coupled NL 104 preamplifiers (Digitimer Ltd, Welwyn, UK). The excitatory postsynaptic potential (fEPSP) was recorded by a 0.1-MΩ tungsten monopolar electrode from the dendritic layer of the stratum radiatum of the CA3 Field following electrical stimulation of the mossy-fiber pathway. The electrical pulses were delivered using a bipolar platinum/iridium electrode (25 lm wire-diameter, at aninter-wire distance of 100 lm, World Precision Instruments, USA). The fEPSP was quantified by 30 % of the maximum slope of its rising phase. Input/output (I/O) curves were constructed by measuring fEPSP slopes responding to the stimulus intensity increasing from 1 to 10 V, with a 0.5-V increment in each slice. Paired-pulse facilitation (PPF) was examined by applying pairs of pulses, which were separated by 20-500 ms intervals.

For induction of long-term potentiation (LTP), we used theta-burst stimulation (TBS) that consisted of 4 pulses at 100 Hz, repeated 3 times with a 200-ms interval. The magnitudes of LTP are expressed as the mean percentage of baseline fEPSP initial slope.

NMDA/AMPA receptor EPSC analysis was performed by patch clamp in the presence of 50 mM picrotoxin. The evoked EPSCs were collected at two holding potentials. At −70 mV, responses were collected and the peak amplitude identified as the AMPA receptor-mediated response. Cells were then voltage clamped at +40 mV, and the amplitude of the evoked EPSC 50 ms post-stimulus was identifed as the NMDAR-mediated response. Six to eight traces were collected at 0.1 Hz for each membrane potential [30–31].

### Cell culture

The human embryonic kidney293 (HEK293) were grown in Dulbecco’s Modified Eagle’s medium (DMEM), supplemented with 10% (v/v) fetal bovine serum and 1% penicillin/streptomycin, in a humidified atmosphere containing 5% CO_2_ incubator at 37 °C. After growing 24 h in plates or flasks, the cells were transfected with the indicated plasmid(s) using Lipofectamine2000 according to the manufacturer’s instructions.

For primary neuron cultures, 18 days embryonic (E18) rat hippocampus were seeded at 30,000-40,000 cells per well on 6-well plates coated with Poly-D-Lysine/Laminin (Bioscience) in neurobasal medium (Invitrogen) supplemented with 2% B27/0.5 mM glutamine/25 mM glutamate. Half the culture medium was changed every 3 days with neurobasal medium supplemented with 2% B27 and 0.5 mM glutamine. All cultures were kept at 37 °C in a humidified 5% CO_2_ containing atmosphere. More than 90% of the cells were neurons after they were cultured for 7 to 17 div; this was verified by positive staining for the neuronal specific markers microtubule-associated protein-2 (MAP2, dendritic marker, Millipore).

### Preparation of nuclear fractionation

The nuclear extracts were prepared using the nuclear extraction kit according to the manufacturer’s instructions (Signosis, Inc., Sunnyvale, CA). Briefly, culture dish was added with Buffer I working reagent and rocked at 200 rpm for 10 min on a shaking platform at 4 °C. The HEK293 cells were collected and centrifuged at 12,000rpm for 5 min at 4 °C. The supernatant was discarded, and the pellets were re-suspended by adding Buffer II working reagents. For tissues, the hippocampal CA3 areas (where virus infected) were rapidly cut into small pieces, added Buffer I working reagent, and homogenized on ice until a single cell suspension observed (by microscope). After spun at 500 g for 5 min at 4 °C, the supernatant was removed, and the cell pellets re-suspended with Buffer I working reagent and rocked at 200 rpm for 10 min on a shaking platform at 4 °C. Then the cells centrifuged at 10,000 rpm for 5 min at 4 °C, and the pellets were re-suspended by adding Buffer II working reagents. Lastly, the cell lysis was shaken at 200 rpm on a platform for two hours at 4 °C. After centrifuged at 12,000 rpm for 5 min at 4 °C, the supernatant (nuclear extract) was collected and stored at −80 °C until use.

### Preparation of insoluble tau

Insoluble tau aggregates were isolated from virus infected hippocampal tissue by a modified procedure. Brain tissues were homogenized in lysis buffer (10 mM Tris-HCl, 150 mM NaCl, 20 mM NaF, 1 mM Na_3_VO_4_, 2 mM EGTA, 0.5% Triton X-100, and 0.1% SDS) with protease inhibitor mixture and centrifuged for 20 min at 13,000 × g. The resulting supernatant designated as soluble tau fraction. The pellet was resuspended in 1% SDS buffer with 10 times ultrasonic and designated as insoluble aggregated tau.

### Western blotting

Equal amounts of protein were separated by 10% sodium dodecyl sulfate-polyacrylamide gel electrophoresis (SDS-PAGE) and transferred onto nitrocellulose membranes. For analysis of STAT1 dimerization, cell lysates were incubated for 20 min with 1 mM DSS, blocked with 0.5 mM NH_4_OH, and used for Western blotting with anti-STAT1 antibody [32]. The membranes were blocked in 5% non-fat milk for 1 h at room temperature and then incubated with primary antibody (Supplementary Table 3) at 4 °C overnight. Then the blots were incubated with IRDye 800CW-conjugated affinity-purified anti-mouse IgG (Rockland) or IRDye 800CW anti-rabbit IgG secondary antibody (Rockland) for 1 h at room temperature. Immunoreactive bands were visualized using Odyssey Infrared Imaging System (Licor Biosciences, Lincoln, NE, USA).

### Reverse transcription and real-time quantitative PCR

Reverse transcription and real-time quantitative PCR were carried out according to manufacturer’s instruction (TaKaRa, Dalian, China). The PCR system contains 3 mM MgCl_2_, 0.5 μM forward and reverse primers, 2 μl SYBR Green PCR master mixes and 2 μl cDNA, and the standards for each gene were prepared using appropriate primers by a conventional PCR. The samples were assayed on a Rotor Gene 300 Real-time Cycler (Corbett Research, Sydney, Australia). The expression level of the interest gene was normalized by the housekeeping gene glyceraldehyde-3-phosphate dehydrogenase (GAPDH), which was not changed by the treatments. PCR primers employed in the present study are as follow: Mmu-GluA1 forward and reverse primers, 5‘-CAATGACCGCTATGAGGG-3’ and 5‘ - AAGGACTGAAACGGCTGA-3’; mmu-GluA2 forward and reverse primers, 5‘ - GTGTCGCCCATCGAAAGTG-3’ and 5‘ - AGTAGGCATACTTCCCTTTGGAT-3’; mmu-Syn1 forward and reverse primers, 5‘-AGGACGAGGTGAAAGC-3’ and 5‘-TCAGTCGGAGAAGAGG-3’ mmu-Syt1 forward and reverse primers, 5‘ - CCATAGCCATAGTTGC-3’ and 5‘-GTTTCAGCATCGTCAT-3’; mmu-GluN1 forward and reverse primers, 5‘ - GTCCACCAGACTAAAGA-3’ and 5‘-TCCCATCATTCCGT-3’ mmu-GluN2A forward and reverse primers, 5‘-CTTTTGAGGACGCC-3’ and 5‘-AAATGAGACCCGATG-3’; mmu-GluN2B forward and reverse primers, 5‘ - GGCTGACTGGCTACG-3’ and 5‘ - CTTGGGCTCAGGGAT-3’; mmu-GAPDH forward primer 5‘-GGAGCGAGATCCCTCCAAAAT-3’ and reverse primer 5‘-GGCTGTTGTCATACTTCTCATGG-3’.

### Transcription factor activating profiling assay

Analysis of the activity of 96 transcription factors (TFs, shown in Supplementary Table 1 and 2) was performed according to the manufacturer’s instructions by using the TF Activation Profiling Plate Array II (Signosis, Inc., Sunnyvale, CA). HEK293 cells were transfected with tau plasmid or its control vector for 48 h, and then, the nuclear protein extracts were prepared according to the manufacturer’s instructions using the Nuclear Extraction Kit (Signosis, Inc., Sunnyvale, CA). A 10 μg sample of nuclear protein extracts was assayed per sample.

### Electrophoresis mobility shift assay (EMSA)

The Non-radioactive EMSA-STAT1 Kit was purchased from Signosis (Sunnyvale, CA). EMSA was performed according the protocol supplied by the manufacturer. Briefly, samples were incubated with a biotinized oligonucleotide probe containing a STAT1 binding site. After incubation, the samples were separated on a non-denaturing polyacrylamide gel and transferred to nylon membranes. The transferred oligonucleotides were immobilized by UV crosslinking. For detection of the oligonucleotides, Streptavidin-HRP was added to the membrane, and the blots were developed by ECL according to manufacturers’ instructions. Competition experiment was performed using excess amounts of unlabeled cold probe containing STAT1 binding site.

### Luciferase reporter assay

Activity of the transcription factors (TFs) was analyzed by using specific luciferase reporter vectors, including pSTAT1-Luc, pCBF-Luc, pPIT1-Luc, pHNF1-Luc, pHOX4C-Luc and pSF1-Luc, respectively (Signosis). These vectors contain a cis-element (DNA binding sequence), a minimal promoter, and a firefly luciferase gene. The activated transcription factors (such as STAT1, CBF, PIT1, HNF1, HOX4C, and SF1) bind to the cis-element and trans-activate the expression of the luciferase gene correlating with the measured luciferase enzyme activity. Therefore, the luciferase activity in this assay represents activation of the transcription factor. Briefly, the HEK293 cells were transfected with tau plasmid or its empty vector control in combination with pSTAT1-Luc (or pCBF-Luc, or pPIT1-Luc, or pHNF1-Luc, or pHOX4C-Luc, or pSF1-Luc) reporter construct and pRL-TK for 48 h. Then the cells were washed with PBS and lysed in 100 μl of the 1×CCLR (Promega). The luciferase activity was measured by following the manufacture’s instruction (Promega). The activity of TFs (i.e. firefly luciferase) was normalized to transfection efficiency by using Renilla luciferase activity (pRL-TK).

To generate luciferase reporter plasmids of GluN1, GluN2A or GluN2B promoter, PCR fragments (Fig. 5) from the mouse genomic DNA were inserted into the BglII and NcoI sites of the pGL3 basic luciferase expression vector (Promega, Madison, WI). Mutation of the pGL3-GluN1/GluN2A/GluN2B luciferase plasmid was introduced using the GeneTailor system (Invitrogen). HEK293 cells were transfected with luciferase reporter plasmids using Lipofectamine Plus (Invitrogen) according to the instructions provided by the manufacturer. To assay the luciferase activity, HEK293 cells were seeded into 24-well plates in DMEM, one day prior to transfection, and co-transfected with pGL3-construct, tau and pRL-TK plasmid. After 24 h, cells were harvested and lysed with 100 μl Passive Lysis Buffer. The cell extracts (20 μl) were used for luciferase activity assay using a Lumat LB9507 luminometer (Berthold) and the Dual Luciferase Reporter (DLR) assay system (Promega).

### Chromatin immunoprecipitation (ChIP) assay

The DNA and protein were cross-linked with 1% formaldehyde for 10 min, washed, and scraped into cold PBS with protease inhibitors. After centrifugation, the cell pellet was re-suspended in buffer (20 mM HEPES, pH 7.9, 25 % glycerol, 420 mM NaCl, 1.5 mM MgCl_2_, 0.2 mM EDTA, protease inhibitors), incubated on ice for 20 min, and centrifuged. The pellet (nucleus) was re-suspended in breaking buffer (50 mM Tris-HCl, pH 8.0, 1 mM EDTA, 150 mM NaCl, 1 % SDS, 2 % Triton X-100, protease inhibitors) and sonicated 5~10 s, and Triton buffer was added (50 mM Tris-HCl, pH 8.0, 1 mM EDTA, 150 mM NaCl, 0.1% Triton X-100). An aliquot was reserved as the input, and the remainder was divided to immunoprecipitate with control mouse IgG (Milipore) or STAT1 (Abcam) antibody followed by incubation with protein G beads. Samples were washed three times in Triton buffer, SDS buffer was added (62.5 mM Tris-HCl, pH6.8, 200 mM NaCl, 2 % SDS, 10 mM DTT, 2 μl of proteinase K (40 mg/ml)), and then samples were vortexed and incubated at 65 °C overnight to reverse cross-linking. DNA was isolated using phenol/chloroform extraction and re-suspended in distilled H_2_O. Primers used for ChIP PCR were as follows: GluN1 forward and reverse primer, 5‘- TAGCATTGGCATTGACCC-3’, 5‘- GCTGGTGCGGTGATGTGA-3’; GluN2A forward and reverse primer, 5‘- TCGGCTTGGACTGATACGTG-3’, 5‘- AGGATAGACTGCCCCTGCAC-3’; GluN2B forward and reverse primer, 5‘- TCTCCACCGTGCTGATGT-3’, 5‘- CTCTCCGAGTCTACCTGTTC-3’. PCR products were analyzed by 2% agarose gel electrophoresis.

### Human brain tissue

Fixed AD (n=6, age 80±7) and the age-matched control (n=5, age 78±5) brains (kind gifts of Dr. Iqbal from the NYS Institute for Basic Research, USA) were cut into sections (40 μm) with a freezing microtome (Leitz,Wetzler, Germany; Kryostat 1720) after anhydration in 25 % sucrose to sinking. The study was approved by the Biospecimen Committee. AD was diagnosed according to the criteria of the Consortium to Establish a Registry for AD and the National Institute on Aging. Diagnoses were confirmed by the presence of amyloid plaques and neurofibrillary tangles in formalin-fixed tissue. Informed consent was obtained from the subjects.

### Immunohistochemistry

In brief, mice were sacrificed by overdose chloral hydrate (1 g/kg) and perfused through aorta with 100 ml 0.9 % NaCl followed by 400 ml phosphate buffer containing 4 % paraformaldehyde. Brains were removed and postfixed in perfusate overnight and then cut into sections (20 μm) with a vibratome (Leica, Nussloch, Germany; S100, TPI). The sections of mice and AD brains were collected consecutively in PBS for immunohistochemistry. Free floating sections were blocked with 0.3% H_2_O_2_ in absolute ethanol for 30 min and nonspecific sites were blocked with bovine serum albumin (BSA) for another 30 min at room temperature. Sections were then incubated overnight at 4 °C with primary antibodies. Immunoreaction was developed using HistostainTM-SP kits and visualized with diaminobenzidine (brown color). Sections were counterstained with hematoxylin, dehydrated through a graded ethanol series, mounted on glass slides, and sealed with glass coverslips. For each primary antibody, 3-5 consecutive sections from each brain were used. The images were observed using a microscope (Olympus BX60, Tokyo, Japan).

### Statistical analysis

All data were collected and analyzed in a blinded manner. Data were expressed as mean ± SD or mean ± SEM and analyzed using SPSS 12.0 statistical software (SPSS Inc. Chicago, IL, USA). Statistical analysis was performed using student’s t-test (two-group comparison), two-way ANOVA or two-way ANOVA followed by Bonferroni’s post hoc test. The level of significance was set at p<0.05. Image pro-plus software was used to calculate fluorescence intensity of STAT1 in nucleus on HEK293 cell staining and immunohistochemical staining intensity of STAT1/pY-STAT1 on human brain slice.

## RESULTS

### Intracellular hTau accumulation induces activation of STAT1

During our studies on tau, we often observe that overexpressing hTau proteins result in changes of other proteins. We thus speculate that hTau accumulation may influence gene expression. To test this, we first conducted a whole-genome mRNA chip screening. Indeed, we detected significant alternations in the level of 520 mRNA molecules (235 increased and 285 decreased) in hTau-expressing cells compared with those expressing the empty vector (Fig. S1), suggesting that intracellular hTau accumulation indeed influences gene transcription. To confirm this point, we measured activity of the transcription factors in nuclear fraction by Transcription Factor Activation Profiling Array (Table S1 and Table S2), in which the activity of 96 transcription factors were monitored using a collection of biotin-labeled DNA probes based on the consensus sequences of individual transcription factor DNA binding sites (Signosis). The results showed that the activity of STAT1 and CBF was significantly increased, while the activity of HNF1, HOX4C, PLAG1, SMUC, VDR, SF-1 and PIT1 decreased remarkably in cells overexpressing hTau (Fig. 1A, B). In protein level measured by Western blotting, only elevation of STAT1 but not CBF was shown in total extracts and the nuclear fraction (Fig. S2).

**Figure 1.**
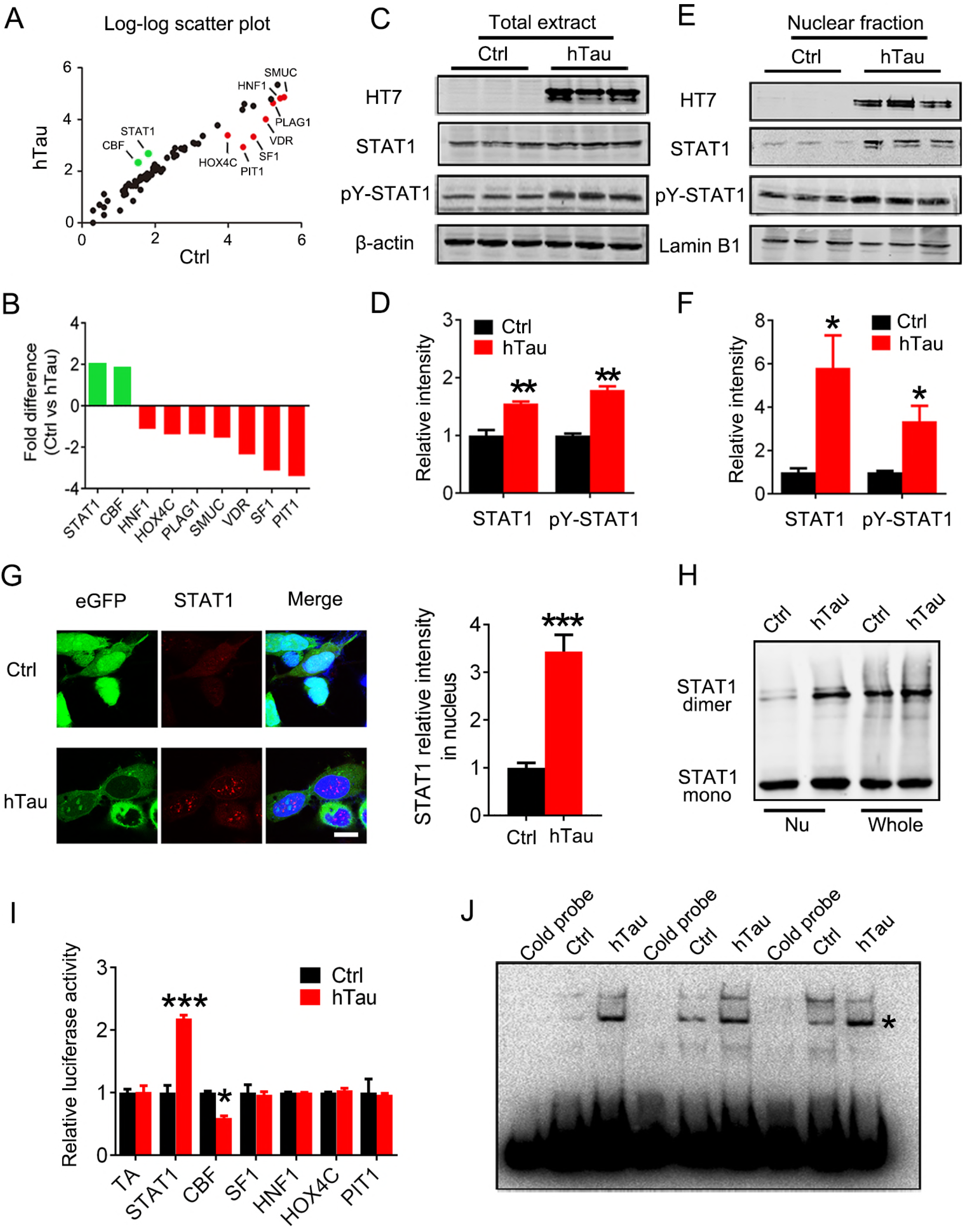
Overexpression of hTau activates STAT1 with an increased nuclear translocation *in vitro*. (A, B) Overexpression of human hTau (hTau) induced significant alterations of 9 transcription factors screened by using Transcription Factors Activation Profiling Plate Array II, in which 96 transcription factors were monitored. The empty vector was transfected as a control (Ctrl). (C-F) Expression of hTau (probed by HT7) increased total and the phosphorylated STAT1 at Tyr701 (pY-STAT1) in cell whole extracts (C, D) and the nuclear fraction (E, F) measured by Western blotting. (G) The representative immunofluorescent images and quantitative analysis show significantly increased STAT1 signal in the nuclear fraction of HEK293 cells with overexpression of hTau compared with the empty vector control (eGFP). Scale bar, 10 μm. (H) Overexpression of hTau most significantly increased STAT1 monomer and dimer formation in nuclear fraction (Nu) measured by Western blotting. (I) Overexpression of hTau increased STAT1 activity in HEK293 cells detected by luciferase assay. (J) Overexpression of hTau increased STAT1-DNA binding activity in HEK293 cells measured by electrophoretic mobility shift assay (EMSA). *, indicates STAT1/DNA complex. Data were presented as mean ±SD of at least three independent experiments (student’s unpaired *t*-test). *, *p*<0.05, **, *p*<0.01, ***, *p*<0.001 vs Ctrl,

Herein, we focused on STAT1 which has been implicated in cognitive functions [33, 34]. We demonstrated that overexpressing hTau remarkably increased the activation-dependent phosphorylation of STAT1 at Tyr701 (pY-STAT1) in both cell lysates (Fig. 1C, D) and the nuclear fraction (Fig. 1E, F) with an enhanced nuclear translocation (Fig. 1G) and dimerization (Fig. 1H) of STAT1 measured by Western blotting and immunofluorescence imaging. Activation of STAT1 by overexpressing hTau was also detected by TFs luciferase assay (Fig. 1I). By EMSA assay using an oligonucleotide probe containing STAT1 binding site, we also found that hTau accumulation increased binding of STAT1 to DNA and this association was disrupted by using cold probe (Fig. 1J). These *in vitro* data indicate that intracellular hTau accumulation induces STAT1 activation.

To test the *in vivo* effects of hTau accumulation on STAT1, we first injected stereotaxically AAV- hTau into the mouse hippocampi and measured the alterations of STAT1 and pY-STAT1 after 1 month. Expression of hTau was confirmed by Western blotting (Fig. 2A), and fluorescent imaging and as well as immunohistochemistry (Fig. S3A). Overexpression of hTau significantly increased total STAT1 and pY-STAT1 in hippocampal extracts and the nuclear fraction (Fig. 2A, B) without changing VDR, PLAG1 and SMUC (Fig. S3B), suggesting a relatively specific effect of hTau on STAT1. Infection of control AAV-eGFP did not activate STAT1 (Fig. S3C). By co-staining of nuclear translocation of STAT1 with NeuN, IBA1 and GFAP, we found that the neuronal staining of STAT1 was most significant (Fig. S4). Elevation of STAT1 and pY-STAT1 was also detected in the hippocampi of 9 m- and 12 m- old hTau transgenic mice (Fig. 2C, D, Fig. S5A). By transfecting Syn-hTau-AAV into the hippocampus, we found that the neuron-specific overexpression of hTau also significantly increased total STAT1 and pY-STAT1 in hippocampal extracts and the nuclear fraction (Fig. S6A, B). In the cortex of AD patients, both total and pY-STAT1 in the nucleus were also significantly increased (Fig. 2E, F). These data provide the *in vivo* and human evidence for the role of tau accumulation in activating STAT1.

**Figure 2.**
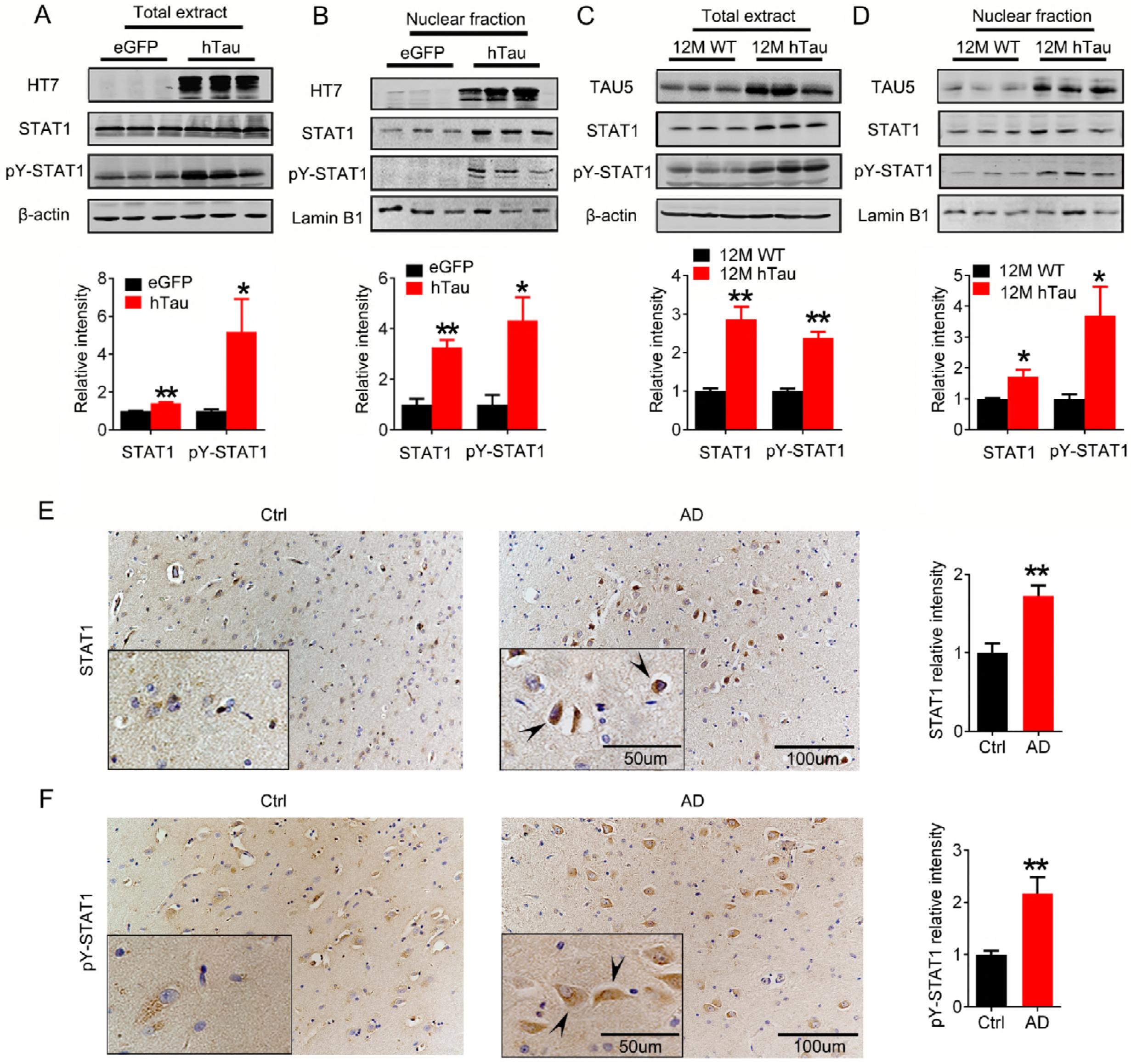
Overexpression of hTau upregulates phosphorylated STAT1 *in vivo*. (A, B) AAV-hTau-eGFP (hTau) or the empty vector AAV-eGFP (eGFP) (1.13×10^13^ v.g./ml) was stereotaxically injected into hippocampal CA3 of 3 m-old C57 mice. After one month, the increased levels of STAT1 and pY-STAT1 in hippocampal total extracts and the nuclear fraction were detected in hTau group by Western blotting. (C, D) The increased STAT1 and pY-STAT1 in hippocampal total extracts and the nuclear fraction of 12 m-old hTau transgenic mice measured by Western blotting. (E, F) The representative images of STAT1 and pY-STAT1 in the brain of AD patients probed by co-immunohistochemical staining and quantitative analysis (hematoxylin stains nuclei, purple; DAB stains the target proteins, brown; n=4-6 slices). Scale bar, 200 μm, or 20 μm for the enlarged. Data were presented as mean ± SD of at least three independent experiments (student’s unpaired *t*-test). *, *p*<0.05, **, *p*<0.01 vs eGFP or wt.

### Downregulating STAT1 rescues hTau-induced memory and synaptic deficits

Previous studies show that accumulation of tau in hippocampal CA3 induces spatial learning and memory deficits in mice [3, 4]. To investigate the role of STAT1 in hTau-induced memory deficits, we co-injected bilaterally AAV-hTau and AAV-Cre into the hippocampal CA3 of STAT1^flox/flox^ mice. The efficiency of AAV-Cre in downregulating STAT1 was confirmed by immunohistochemistry and Western blotting (Fig. 3A, B). By MWM test, we observed that STAT1 knockdown could efficiently rescue the hTau-induced spatial learning impairments shown by the decreased escape latency at days 4 and 5 during the 5-days training (Fig. 3C). In memory test measured at day 6 by removed the escape platform, the mice with STAT1 knockdown showed less average latency to reach the previous target quadrant (Fig. 3D), more frequent crosses in the platform area (Fig. 3E) and more time stayed in the platform quadrant (Fig. 3F) than the control mice. No significant difference in swimming speed was seen among the three groups (Fig. 3G), which excluded motor deficits. By fear conditioning test, we also observed that STAT1 knockdown improved long-term memory shown by an increased freezing time during memory test in human hTau-expressing mice (Fig. 3H). These data demonstrate that downregulating STAT1 in hippocampus can efficiently rescue hTau-induced learning and memory impairments.

**Figure 3.**
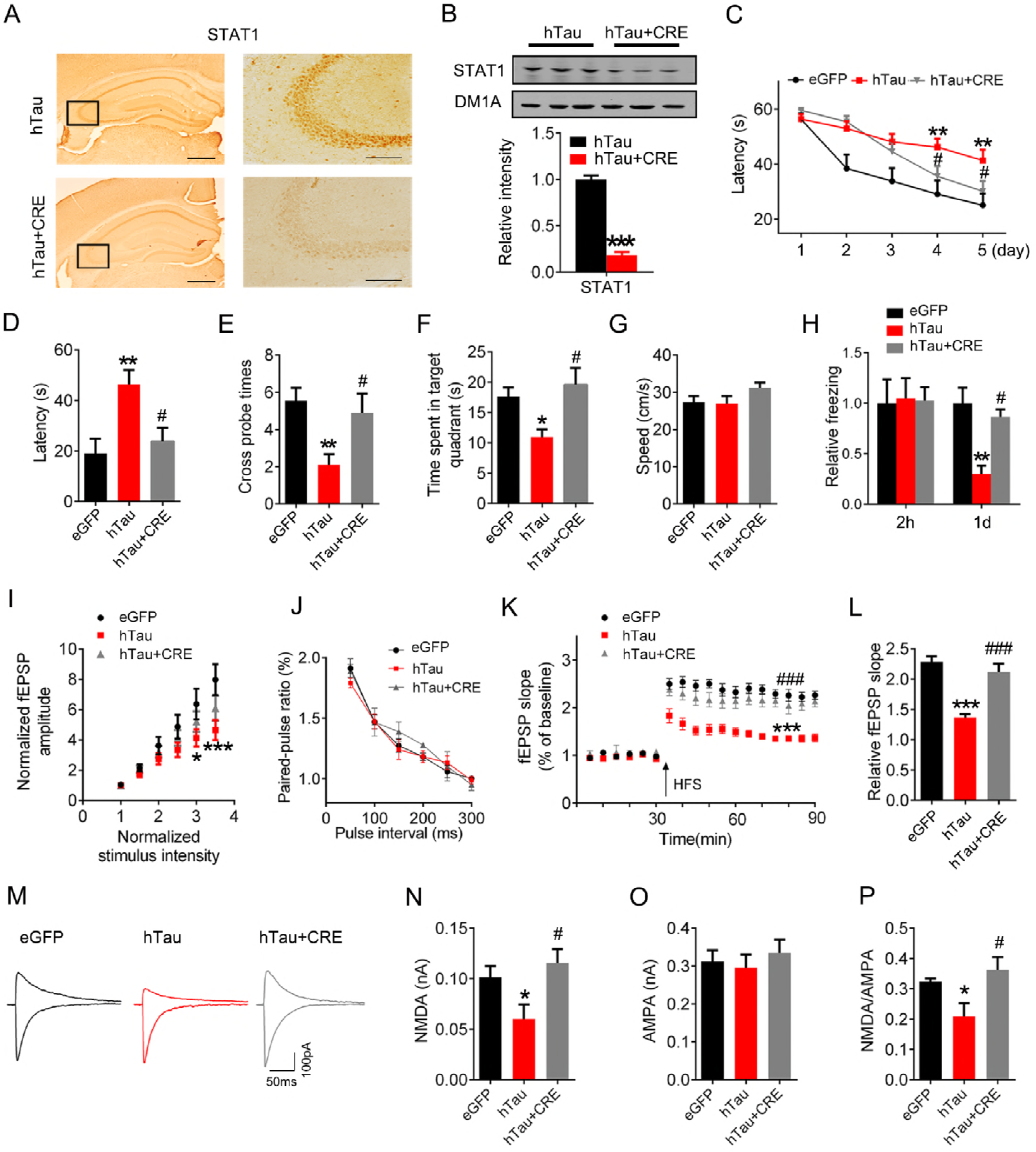
Downregulating STAT1 ameliorates hTau-induced cognitive and synaptic impairments. (A, B) AAV-Cre (5×10^12^ v.g./ml) mixed with AAV-hTau or AAV-eGFP (1.13×10^13^ v.g./ml) were stereotaxically injected into the hippocampal CA3 of 3 m-old STAT1^flox/flox^ mice. One month later, downregulation of STAT1 was confirmed by Western blotting and immunohistochemical staining. Scale bar, 200 μm (left); 100 μm (right); n=3 each group. (C) Downregulation of STAT1 ameliorated hTau-induced spatial learning deficit shown by the decreased escape latency during 5 consecutive days training in Morris water maze (MWM) test (n=9-11 for each group). (D-G) Downregulation of STAT1 ameliorated hTau-induced spatial memory deficit shown by the decreased latency to reach the platform quadrant (D), the increased crossing time in the platform site (E) and time spent in the target quadrant (F) measured at day 6 by removed the platform in MWM test; no motor dysfunction was seen (G) (n=9-11 for each group). (H) Downregulation of STAT1 ameliorated hTau-induced contextual memory deficits measured at 24 h during fear conditioning test (n=8 each group). (I-L) Downregulation of STAT1 restored slopes of field excitatory postsynaptic potential (fEPSP) with no influence on paired-pulse ratio (PPR) recorded in hippocampal CA3 of hTau or STAT1 knockdown mice (n=5 slices from 4 mice for each group). (M-P) One month after the virus infection, whole cell patch clamp was used to measure the function of NMDA (at +40 mV) and AMPA (at −70 mV) receptors on acute brain slices (400 μm). The insets show representative sample traces of EPSCs in virus infected neurons (M). The reduced NMDA and unchanged AMPA currents with a reduced NMDA/AMPA ratio were seen in hTau infected neurons, while knockdown of STAT1 restored the hTau-induced NMDA currents (N-P). (n = 12 neurons from 4 animals for eGFP group; n = 11 neurons from 4 animals for hTau group; n = 13 neurons from 4 animals for hTau+CRE group). Data were presented as mean ± s.e.m for C-H and mean ± SD for others (two-way repeated measures analysis of variance (ANOVA) followed by Bonferroni’ s post hoc test for C, two-way analysis of variance (ANOVA) followed by Bonferroni’ s post hoc test for I-K, one-way analysis of variance (ANOVA) followed by Bonferroni’ s post hoc test for others). *, *p*<0.05, **, *p*<0.01, ***, *p*<0.001 vs eGFP; #, *p*<0.05, ###, *p*<0.001 vs hTau.

Synaptic plasticity is the precondition of learning and memory, therefore we studied how hTau accumulation or with simultaneous STAT1 knockdown affects synaptic functions on the acute brain slices. Using a paired-pulse protocol to determine the paired-pulse ratios (PPR) of the fEPSP at mossy fiber-CA3 circuit, we did not find significant difference between AAV-hTau and AAV-GFP injected mice (Fig. 3J), indicating no significant presynaptic dysfunction. On the other hand, the fEPSP slope was reduced in hTau-expression slices compared with AAV-GFP controls, and downregulating STAT1 substantially attenuated the hTau-induced suppression of LTP (Fig. 3I, K, L). These data indicate that hTau suppresses LTP by preferentially affecting postsynaptic machineries and knockdown STAT1 can rescue the hTau-induced suppression of synaptic transmission.

Using whole-cell patch clamp recording, we measured NMDA and AMPA receptor-mediated synaptic responses at DG-CA3 synapses on acute hippocampal slices. AMPA receptor-mediated responses had no change, while the NMDAR-mediated responses were significantly decreased with a decreased ratio of NMDA/AMPA in hTau-overexpressing mice, and downregulating STAT1 substantially attenuated the hTau-induced suppression of NMDAR currents (Fig. 3M-P). These data provide functional evidence supporting NMDAR impairment by hTau accumulation in CA3 neurons.

### STAT1 suppresses NMDAR expression *via* binding to the specific domain of the promoter

To explore how hTau-induced STAT1 elevation affects synaptic function, we measured the level of synapse-related proteins. The results showed that hTau accumulation in mice or overexpression WT-STAT1 in primary hippocampal neurons decreased the protein and mRNA levels of postsynaptic proteins N-methyl-D-aspartate receptors (NMDARs) type 1 (GluN1), GluN2A and GluN2B, while knockdown STAT1 by AAV-Cre substantially restored the protein and mRNA levels of the NMDARs measured respectively by Western blotting (Fig. 4A, B, D, E; Fig. S7A, B), RT-PCR (Fig. 4C, F; Fig. S7C), and immunohistochemical staining (Fig. S7D, E). On the other hand, overexpressing hTau with or without STAT1 knockdown did not significantly affect the protein levels of presynaptic proteins synapsin1 (Syn1) and synaptotagmin1 (Syt1), or postsynaptic proteins AMPA receptor subunits GluA1 and GluA2 (Fig.4A-F). These data suggest that STAT1 elevation mediates the hTau-induced suppression of NMDAR expression. We also found that NMDAR protein levels decreased in the 9 m- and 12 m-old hTau transgenic mice compared with the wildtype littermates (Fig. S5B). By transfecting Syn-hTau-AAV into the hippocampus, we found that the Syn-specific neuronal overexpression of hTau also decreased NMDAR levels (Fig. S6C), as seen in the pan-neuronal overexpression of hTau (Fig. 4A, 4B). To explore how STAT1 suppresses the expression of NMDARs, we screened potential binding sites of STAT1 in the promoter regions of GluN1, GluN2A and GluN2B in a transcription factor database [35]. We found 2 conserved GAS promoter elements for STAT1 binding in the promoter regions of GluN1 and GluN2B, and 4 GAS promoter elements in GluN2A (Fig. 5C, E, G). Further studies by chromatin immunoprecipitation (CHIP) assay demonstrated that overexpression of hTau in hippocampus remarkably increased binding of STAT1 to the promoters of GluN1, GluN2A and GluN2B genes (Fig. 5A), and upregulating wildtype STAT1 inhibited transcription activity of the NMDARs (Fig. 5B). These data together demonstrate that STAT1 activation can suppress NMDAR expression by direct binding to the promoter.

**Figure 4.**
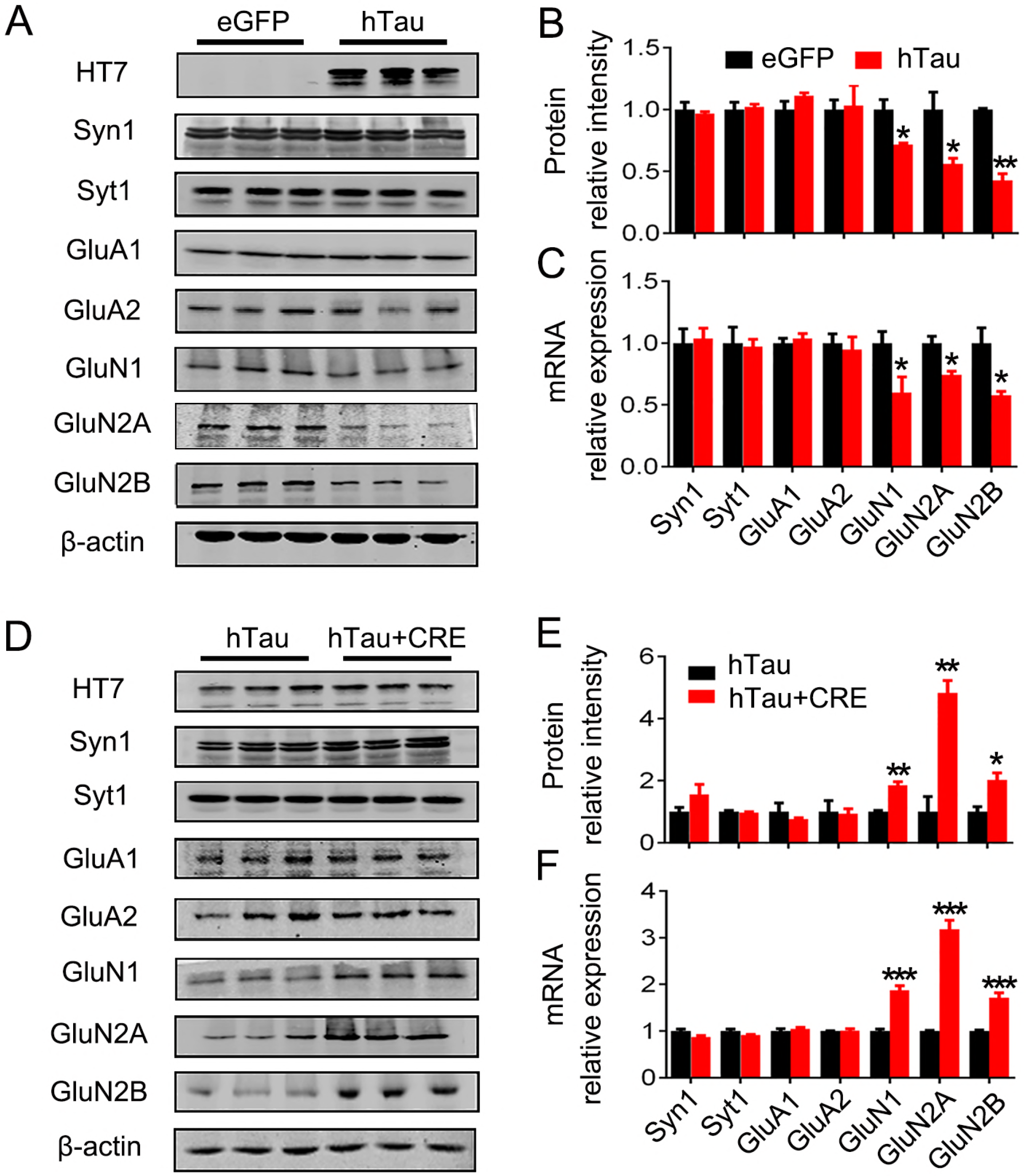
Overexpressing hTau suppresses expression of NMDARs by upregulating STAT1 in mice. (A-C) Overexpression of AAV-hTau decreased the protein and mRNA levels of GluN1, GluN2A and GluN2B detected by Western blotting and qRT-PCR in the hippocampal CA3 of C57 mice, compared with the AAV-eGFP vector control. Data were presented as mean ± SD (n=3-4; student’s unpaired t-test). (D-F) Simultaneous downregulation of STAT1 by infusing AAV-Cre in hippocampal CA3 of STAT1^flox/flox^ mice abolished the hTau-induced inhibition in expression of NMDAR protein and mRNA. Data were presented as mean ± SD (n=3-4; student’s unpaired *t*-test). *, *p*<0.05, **, *p*<0.01, ***, *p*<0.001 vs eGFP or hTau.

**Figure 5.**
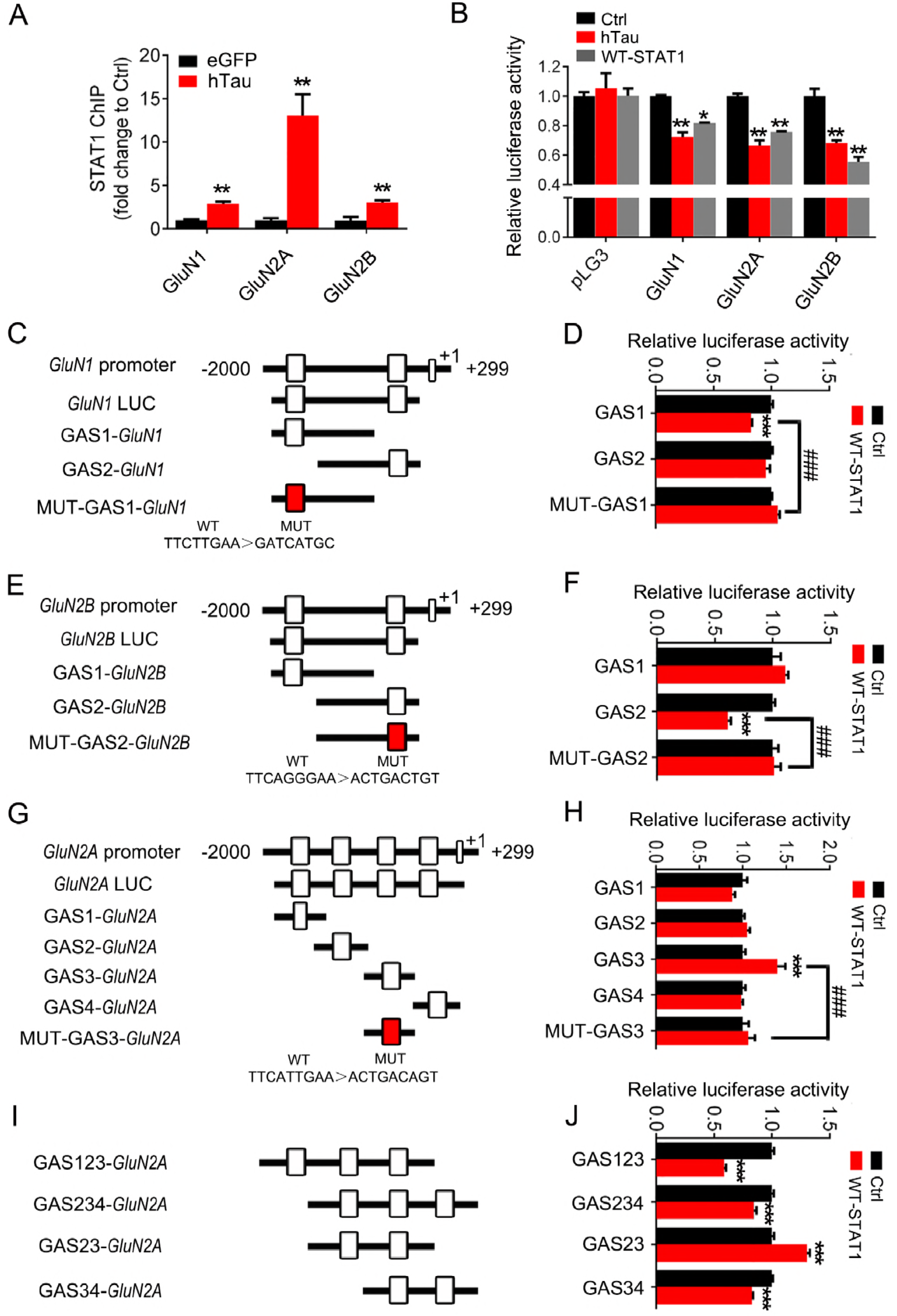
Overexpressing hTau increases binding of STAT1 to *NMDAR* promoters and inhibits the expression of NMDARs. (A) Overexpression of AAV-hTau increased binding of STAT1 to the promoter regions of *GluN1*, *GluN2A* and *GluN2B* gene in hippocampal CA3 extracts measured by chromatin immunoprecipitation assay (CHIP). (B) Overexpression of hTau or wildtype STAT1 (WT-STAT1) inhibits the transcription activity of NMDARs compared with the empty vector control (Ctrl) measured by luciferase activity assay in HEK293 cells. (C-J) Diagrams show the predicted GAS promoter element (GASs) for STAT1 in the promoter (−2000-+299bp) of GluN1 (C), GluN2B (E) and GluN2A (G, I). The GASs or the mutant (MUT) plasmids were co-transfected respectively with WT-STAT1 or its empty vector (Ctrl) into HEK293 cells for 24 h, and then the luciferase activity was measured (right panels). n=4 for each group. Data were presented as mean ± SD (two-way analysis of variance (ANOVA) followed by Bonferroni’ s post hoc test). *, *p*<0.05, **, *p*<0.01, ***, *p*<0.001 vs Ctrl; #, *p*<0.05, ###, *p*<0.001 vs wildtype reporters.

To clarify the specific GAS promoter element of GluN1, GluN2A or GluN2B genes for STAT1, we constructed luciferase reporters containing various GAS elements on the NMDAR promoters (Fig. 5C-H). After co-transfection of specific GAS element reporters with STAT1 into HEK293 cells, we found that co-expression of STAT1 with GAS1 on GluN1 (Fig. 5C, D) or GAS2 on GluN2B (Fig. 5E, F) induced inhibition of luciferase activity, while the luciferase activity of GAS2 on GluN1 and GAS1 on GluN2B was not changed by STAT1 (Fig. 5D, F). Furthermore, expression of mutant GAS1 on GluN1or GAS2 on GluN2B abolished STAT1-induced inhibition of luciferase activity (Fig. 5D, F). These data suggest that STAT1 inhibits GluN1 and GluN2B expression by binding to GAS1 (GluN1) and GAS2 (GluN2B) elements, respectively.

In case of GluN2A that has 4 GAS elements, we found that co-expression of STAT1 with GAS1, 2 or 4 elements did not change luciferase activity (Fig. 5G, H); but co-expression of STAT1 with GAS3 element induced transcriptional activation and that was abolished by GAS3 mutant (Fig. 5H). To clarify these conflict results, we did a random assortment study of the reporters. The results showed that co-expression of GAS1-3, GAS2-4 and GAS3-4 elements with STAT1 induced inhibition of luciferase activity, while co-expression of GAS2-3 elements increased the luciferase activity (Fig. 5I-J). These data suggest that a multi-GASs-dependent binding of STAT1 may be involved on GluN2A subunit.

We also measured laminin β1 (LB1) that is involved in Aβ-induced suppression of NMDAR expression [33]. No significant change was detected after overexpressing hTau (Fig. S8), suggesting that hTau induces synapse impairment with distinct mechanisms from Aβ.

### JAK2 activation mediates hTau-induced STAT1 upregulation

Phosphorylation of STAT1 is critical for its nuclear translocation and the activation [36]. To further explore the upstream factors mediating hTau-induced STAT1 activation, we screened protein kinases that can phosphorylate STAT1 [37, 38]. Among various kinases, JAK2, JNK and ERK were activated by overexpressing hTau (Fig. 6A, B), while only simultaneous inhibition of JAK2 by JAK2 inhibitor TG-101348 (JAK2I) or JAK2 siRNA but not JNK or ERK abolished hTau-induced STAT1 hyperphosphorylation at pY701 in both total cell extracts and the nuclear fraction (Fig. 6C-J, Fig. S9). JAK2 activation was also detected in hippocampus of 12 m-old hTau transgenic mice and CMV-hTau-AAV or Syn-hTau-AAV infused C57 mice (Fig. 6K, L and Fig. S6A). These data demonstrate that hTau accumulation upregulates STAT1 activity by activating JAK2.

**Figure 6.**
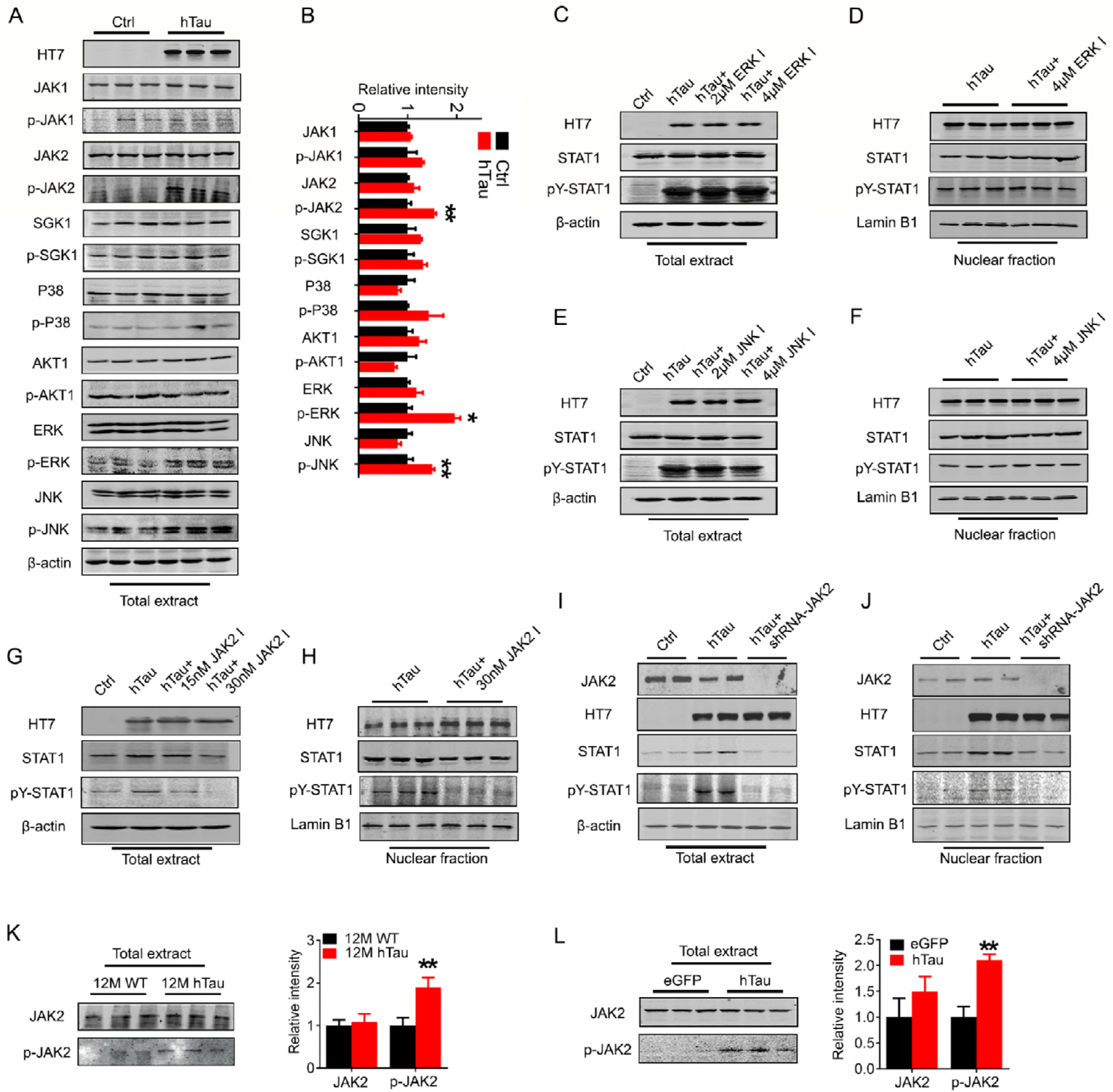
JAK2 activation mediates hTau-induced STAT1 activation. (A, B) Overexpression of hTau in HEK293 cells for 48 h increased the activity-dependent phosphorylation of JAK2, JNK1 and ERK1 compared with the empty vector control (Ctrl) measured by Western blotting (n=3). (C-F) Pharmacological inhibition of ERK1 (C, D) or JNK1 (E, F) for 24 h did not significantly affect the hTau-induced STAT1 phosphorylation at pY-STAT1 (Tyr701) in total extracts (C, E) and the nuclear fraction (D, F) measured by Western blotting (n=3). The alteration of pS-STAT1 (Ser727) confirms the efficacy of JNK1 inhibitors. (G-J) Pharmacological inhibition of JAK2 (G, H) or knockdown JAK2 by siRNA (I, J) abolished hTau-induced STAT1 phosphorylation at Tyr701 in total extracts (G, I) and the nuclear fraction (H, J) (n=3). (K, L) The phosphorylated JAK2 level increased in the hippocampus of 12 m-old hTau transgenic mice (K), and the hippocampus of C57 mice infected with AAV-hTau (1.13×10^13^ v.g./ml) (L). Data were presented as mean ± SD (student’s unpaired t-test); *, *p*<0.05, **, *p*<0.01 vs Ctrl, eGFP or WT.

### Blocking STAT1 activation rescues hTau-induced synapse and memory impairments

To verify the role of STAT1 phosphorylation in regulating expression of NMDARs and the cognitive ability, we constructed non-phosphorylation STAT1 dominant negative mutant (Y701F-STAT1) AAV virus and co-infused the mutant virus with AAV-hTau into the hippocampal CA3 of 3 m-old C57 mice for one month (Fig. 7A). We found that co-expression of dominant negative Y701F-STAT1 attenuated hTau-induced learning and memory deficits (Fig. 7B-F) with attenuation of LTP suppression (Fig. 7G, H) and restoration of GluN1, GluN2A and GluN2B protein and mRNA levels (Fig. 7I-K). These data reveal that phosphorylation of STAT1 at Tyr701 indeed plays a critical role in hTau-induced synapse and memory impairments.

**Figure 7.**
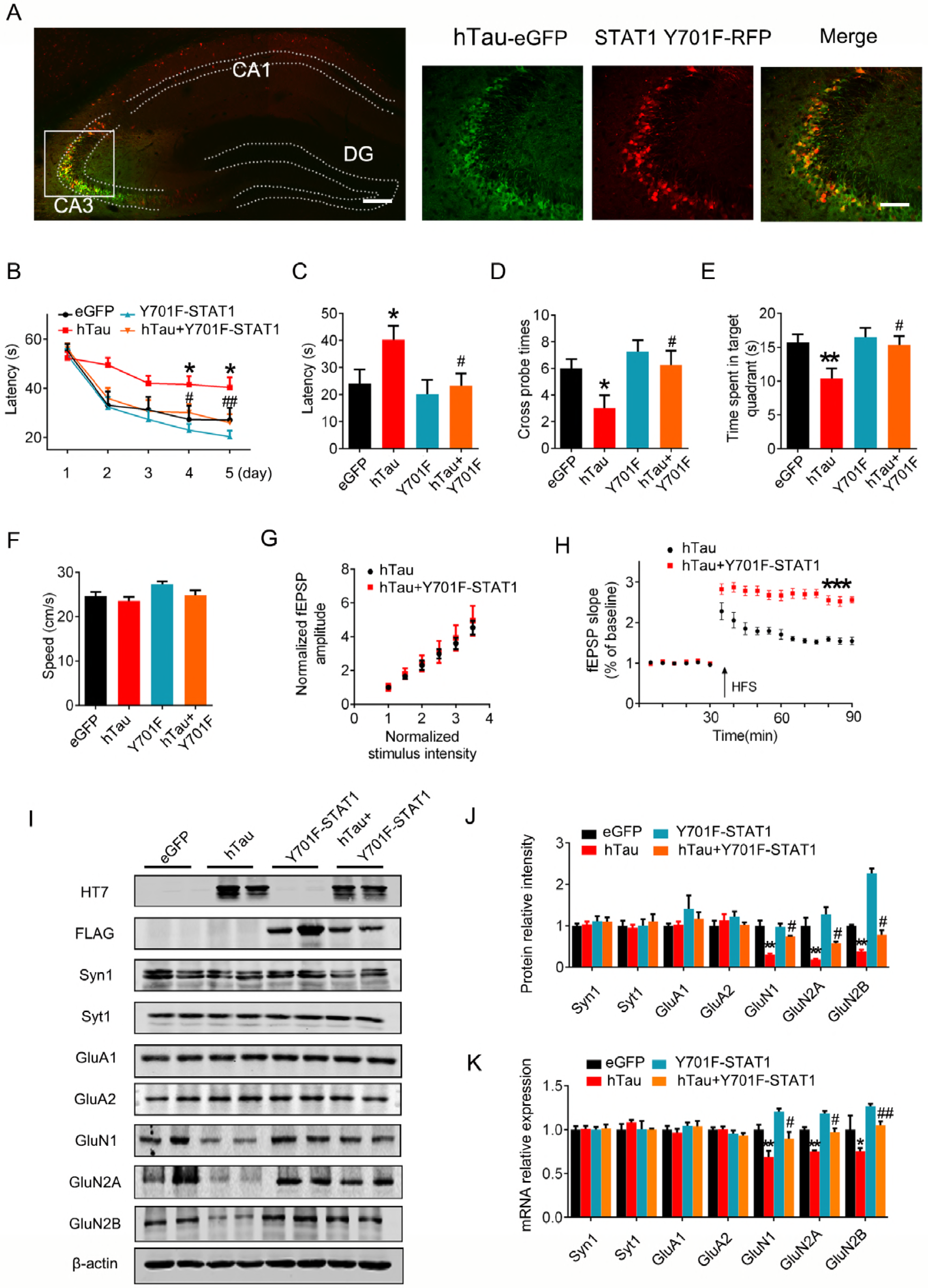
Blocking STAT1 activation rescues hTau-induced synapse and memory impairments. (A-E) AAV-eGFP (eGFP) or AAV-hTau-eGFP (hTau) (1.13×10^13^ v.g./ml) or AAV-Y701F-STAT1 (5×10^12^ v.g./ml) or AAV-Y701F-STAT1 (5×10^12^ v.g./ml) plus hTau was stereotaxically injected into hippocampal CA3 of 3 m-old C57 mice. After one month, learning and memory were detected by MWM test. (A) The representative fluorescence image confirms expression of AAV-hTau and AAV-Y701F-STAT1. (B) Overexpression of Y701F-STAT1 mitigated hTau-induced spatial learning deficits shown by the decreased escape latency during water maze training. Data were presented as mean ± s.e.m (n=7-10 each group, two-way analysis of ANOVA, Bonferroni’s post hoc test). (C-E) Overexpression of Y701F-STAT1 mitigated hTau-induced spatial memory impairment shown by the decreased latency to reach the platform (C), the increased crossing time in the platform site (D) and time spent in the target quadrant (E) measured at day 6 by removed the platform (n=7-10 each group). Data were presented as mean ± s.e.m (two-way ANOVA, Bonferroni’s post hoc test). (F) Expression of Y701F-STAT1 did not change the swimming speed of the mice in water maze task. (G) Simultaneous expression of Y701F-STAT1 did not induced any further change on basal synaptic transmission (I/O curve) compared with expression of hTau alone, recorded in hippocampal CA3 (n=5 slices from 4 mice for each group). (H) Simultaneous expression of Y701F-STAT1 abolished the hTau-induced inhibition of field excitatory postsynaptic potential (fEPSP), recorded in hippocampal CA3 of C57 mice (student’s unpaired t-test, n=5 slices from 4 mice for each group). (I-K) Simultaneous expression of Y701F-STAT1 rescued the hTau-induced suppression of NMDARs protein (I, J) and mRNA (K) expression measured by Western blotting and qRT-PCR in hippocampal CA3 of C57 mice. Data were presented as mean ± s.e.m for B-F and mean ± SD for others (two-way repeated measures analysis of variance (ANOVA) followed by Bonferroni’ s post hoc test for B, two-way analysis of variance (ANOVA) followed by Bonferroni’ s post hoc test for others, n=3 each group). *, *p*<0.05; **, *p*<0.01 vs eGFP or hTau; #, *p*<0.05, ##, *p*<0.01 vs hTau.

To explore whether STAT1 knockdown affects tau phosphorylation and aggregation, we co-infused STAT1 dominant negative mutant (AAV-Y701F-STAT1) with AAV-hTau into the hippocampal CA3 of 3 m-old C57 mice. After one month, the hTau level in soluble and insoluble fractions of hippocampal CA3 was measured. Reduction of p-hTau (pS214, pT231 and pS404) was shown in soluble and insoluble fractions, and total tau (Tau-5) protein decreased in insoluble fractions (Fig. S10). These data suggest that downregulating STAT1 could attenuate hTau-toxicities by reducing tau hyperphosphorylation and the pathological aggregation.

## DISCUSSION

Tau accumulation forming neurofibrillary tangles is hallmark of AD pathologies, but how tau accumulation induces synapse and memory impairment is elusive. By overexpressing hTau to mimic intraneuronal tau accumulation as seen in the sporadic AD cases, we show that hTau accumulation activates JAK2 to phosphorylate and activate STAT1. Upregulation of STAT1 subsequently inhibits expression of GluN1, GluN2A and GluN2B by binding to their specific promoter elements, which results in synaptic dysfunction and memory deficit. We also demonstrate that knockdown STAT1 by AAV-Cre in STAT1^flox/flox^ mice or by overexpressing dominant negative AAV-Y701F-STAT1 efficiently rescues hTau-induced suppression of NMDAR expression with attenuation of synaptic functions and memory performance. These findings reveal that intracellular accumulation of hTau causes memory deterioration through JAK2/STAT1-induced suppression of NMDARs expression, which discloses a novel mechanism for tau-related synapse and memory impairments (Fig. S11).

The mammalian STAT family is consisted of seven members, i.e., STAT1, 2, 3 and 4, STAT5a, STAT5b and STAT6 [39], among them, STAT1, 3, 5 and 6 are differentially expressed in the brain [40]. Serve as transcription factors, the activity of STATs is regulated by phosphorylation [41, 42], and phosphorylation of STAT1 at Tyr701 stimulates its dimerization, nuclear translocation, DNA binding, and activation [36]. By using multiple measures including phosphorylation, dimerization, EMSA and luciferase activity assay, we provide strong evidence showing that hTau accumulation can activate STAT1. Furthermore, expression of un-phosphorylable dominant negative Y701F-STAT1 attenuates hTau-induced suppression of synaptic plasticity, which confirms a critical role of Tyr701-phosphorylation in regulating STAT1 activity. In addition to STAT1, we also detected upregulation of CBF and downregulation of seven Tfs in HEK293 cells after overexpressing hTau by transcription factor activating profiling assay. In the following studies on HEK293 and mouse brain with overexpression of hTau, some of the results were not recapitulated by Western blotting. This discrepancy can be caused by different measures (activity *versus* protein level), the experimental methods and the materials used. By website prediction of the transcription factor binding sites (http://gene-regulation.com/pub/programs/alibaba2/index.html), we also found that in addition to STAT1, the other Tfs, such as HNF1, HOX4C, and PIT1, also have potential NMDAR binding element. Therefore, we measured whether overexpressing hTau affect the activity of HNF1, HOX4C, and PIT1, but no significant change was shown. These data suggest a relatively specific and significant effect of hTau on STAT1, and consequently the role of STAT1 on NMDARs.

Several tyrosine kinases, such as ERK1, JNK1, p38 kinase, MEK1, MSK1 and the JAK kinases, are involved in STAT1 phosphorylation [37, 38]. Among them, we observed that JAK2, JNK and ERK were activated upon intracellular hTau accumulation. However, only simultaneous inhibition of JAK2 but not JNK and ERK abolished the hTau-induced STAT1 phosphorylation, which suggests a critical role of JAK2 activation in hTau-induced STAT1 activation. The JAK/STAT pathway is involved in many pathophysiological processes including cell survival, proliferation, differentiation, development and inflammation. Recent studies show that overexpression of STAT1 impairs water maze performance in mice [33, 34]. STAT1 can bind to the promoter of extracellular matrix protein laminin β1 (LB1), by which it downregulates the expression of GluN1 and GluN2B in Aβ treatment [33]. These data suggest an indirect role of STAT1 in regulating NMDARs via LB1. In the present study, we find that STAT1 can directly bind to the special GAS elements on NMDAR promoters, and thus directly blocks the expression of NMDAR subunits. These data not only reveal novel mechanism underlying the STAT1-regulation on synaptic function, but also provide potential strategy for intervention.

To identify the exact binding element(s) of STAT1 on NMDAR promoters, we constructed GAS promoter elements (GASs) in NMDAR promoter regions for luciferase activity assay. We observed that GAS1 in GluN1 promoter and GAS2 in GluN2B promoter were required for STAT1 negative regulation of the genes expression. However, STAT1 increased luciferase activity of GAS3-containing constructs in GluN2A promoter, which is inconsistent with the reduced mRNA and protein levels of GluN2A by STAT1. Our further studies reveal that STAT1 negative regulation of GluN2A promoter needs concomitant effect of GAS3 with GAS1 or GAS4, suggesting that STAT1-dependent suppression of GluN2A requires a distal promoter region containing multiple GAS elements. This phenomenon was also seen in IFNγ-mediated transcriptional suppression of the perlecan gene [43]. It is well known that STAT1 plays an important role in immune response [44], and inhibition of neuroinflammation ameliorates learning and memory deficits in AD animal models [45–47]. Whether and how inflammation may be involved in hTau-induced memory deficits deserve further investigation.

Taken together, we find here that intracellular accumulation of hTau suppresses NMDAR expression by activating JAK2/STAT1 signaling pathway, and thus induces synaptic and memory impairments. Downregulating STAT1 or blocking STAT1 activation efficiently rescues the hTau-induced synaptic dysfunction and memory impairment in mice.

## ACKNOWLEDGMENTS

We thank Dr. Xiao-Yuan Li of Institute of Biomedical Sciences, Academia Sinica, Taiwan for the kind gift of STAT1 plasmids. This work was supported in parts by Natural Science Foundation of China (81471303, 31730035 and 81801062), and by Ministry of Science and Technology of China (2016YFC1305800).

## AUTHOR CONTRIBUTIONS

J.Z.W. and G.P.L. conceived the project, designed the experiments, and wrote the manuscript. X.G.L and X.Y.H designed and performed most of the experiments. Y.L.W, S.J.Z., and D.S.S. performed electrophysiological experiments. Q.F., J.W.Y. and Y.G. prepared primary neurons. H.L.L. performed the immunohistochemical experiments. J.F.Z., X.C.L., Y.C.L., D.K., and Q.W. assisted with *in vivo* and *in vitro* experiments. K.Y. assisted with data analysis and interpretation and critically read the manuscript.

## CONFLICTS OF INTEREST

All authors disclose: (a) no actual or potential conflicts of interest including any financial, personal or other relationships with other people or organizations within three years of beginning the work submitted that could inappropriately influence (bias) their work. (b) When applicable, provide statements verifying that appropriate approval and procedures were used concerning animals.

